# Evidence for Transcriptome-wide RNA Editing Among *Sus scrofa* PRE-1 SINE Elements

**DOI:** 10.1101/096321

**Authors:** Scott A. Funkhouser, Juan P. Steibel, Ronald O. Bates, Nancy E. Raney, Darius Schenk, Catherine W. Ernst

## Abstract

**Background:** RNA editing by ADAR (adenosine deaminase acting on RNA) proteins is a form of transcriptional regulation that is widespread among humans and other primates. Based on high-throughput scans used to identify putative RNA editing sites, ADAR appears to catalyze a substantial number of adenosine to inosine transitions within repetitive regions of the primate transcriptome, thereby dramatically enhancing genetic variation beyond what is encoded in the genome.

**Results:** Here, we demonstrate the editing potential of the pig transcriptome by utilizing DNA and RNA sequence data from the same pig. We identified a total of 8550 mismatches between DNA and RNA sequences across three tissues, with 75% of these exhibiting an A-to-G (DNA to RNA) discrepancy, indicative of a canonical ADAR-catalyzed RNA editing event. When we consider only mismatches within repetitive regions of the genome, the A-to-G percentage increases to 94%, with the majority of these located within the swine specific SINE retrotransposon PRE-1. We also observe evidence of A-to-G editing within coding regions that were previously verified in primates.

**Conclusions:** Thus, our high-throughput evidence suggests that pervasive RNA editing by ADAR can exist outside of the primate lineage to dramatically enhance genetic variation in pigs.

## Background

Eukaryotes are known for relatively complex mechanisms used to regulate gene expression. One such mechanism, RNA editing, enables the cell to alter sequences of RNA transcripts [1] such that they are no longer forced to match the “hard-wired” genome sequence. High throughput methods for studying targets of this mechanism transcriptome-wide have been applied to primate studies, where evidence for massive amounts of ADAR (adenosine deaminase acting on RNA) catalyzed A-to-I RNA editing has been discovered, preferentially within SINE retrotransposons such as the primate Alu [2–8]. Such work has yet to be performed with pig transcriptomes using the latest sequencing technology. Although little is known about pig SINE elements compared to those in primates, key features of the pig-specific PRE-1 retrotransposon make pigs an intriguing model to further elucidate transcriptome-wide patterns of ADAR targets.

ADAR can only catalyze A-to-I editing within dsRNA. The high editibility of the primate specific Alu element is attributed to its capacity to induce dsRNA; these elements have a high copy number, are short, relatively undiverged from one another, and tend to cluster in gene rich regions of the genome [9]. When appearing as tandem and inverted pairs within the same transcribed region, these properties facilitate intra-molecular dsRNA formation that serve as ADAR targets [2, 10]. Comparatively, the pig PRE-1 element possesses many of these same properties that are believed to contribute to dsRNA formation within the transcriptome. Notably, PRE-1 has the 3^rd^ highest copy number of any SINE cataloged on SINEBase [11].

Since Alu elements are generally found within and near genes, ADAR editing in humans preferentially targets non-coding regions of many genes such as introns, UTRs and upstream and downstream gene proximal regions. ADAR editing of these regions is thought to be a key component of RNA processing via mechanisms that include Alu exonization [12] and RNAi pathway alteration [13]. By demonstrating that RNA editing in pigs generally targets SINE elements within non-coding regions of genes, this would suggest that RNA processing by way of ADAR editing of SINE elements predated the emergence of primate and pig-specific retrotransposons. Rarely, ADAR editing occurs within coding regions to alter amino acid sequences [14]. This type of editing is particularly mysterious in that its pattern is less traceable than non-coding editing, but is nevertheless site-specific and required for the function of essential protein coding genes such as *GluR-B* in mice [15]. Therefore, in addition to the regulation of transcripts by way of editing non-coding SINE elements, editing of coding regions is an essential form of transcriptional regulation in mice, with the extent of its conservation across Mammalia yet to be fully determined.

Here, we demonstrate the pig’s capacity for RNA editing. By studying this process in a relatively distant species to human with a distinct repetitive element repertoire, we want to determine if RNA editing patterns seen in Alu bearing genomes can likewise be observed in pigs. RNA editing detection was done by analyzing a single pig using whole genome sequencing data and RNA sequencing data from liver, subcutaneous fat, and *longissimus dorsi* muscle. Based on previous studies done in primates, a bioinformatic strategy was used to find A-to-I (observed as A-to-G) DNA to RNA mismatches that give evidence of ADAR catalyzed RNA editing events.

## Results and discussion

### DNA and RNA sequencing

To provide the materials needed for a transcriptome-wide survey of RNA editing candidates, genomic DNA as well as total RNA from liver, subcutaneous fat, and *longissimus dorsi* (LD) muscle were purified from samples obtained from a single animal, similar to another single-animal editome study [8]. Sequencing was done using the Illumina HiSeq 2500 to generate 150x2 paired end reads from genomic DNA, with PolyA RNA sequencing used to generate cDNA reads in the same format. Roughly 250M pass-filter genomic DNA reads were generated with an average overall alignment rate of 89% to the *Sus scrofa* reference genome sequence (*Sus scrofa* 10.2.69). An average of 106M pass-filter strand specific cDNA reads were obtained from each tissue, with an average overall alignment rate of 76%.

### Identification of candidate RNA editing events

To scan the transcriptome for possible RNA editing sites, we utilized a custom pipeline influenced by previous studies done in human cell lines and primates [16, 8]. Prior to alignment, in order to avoid utilizing bases with relatively poor base qualities at the ends of reads, raw genomic DNA and cDNA sequencing reads were trimmed for base quality at their 3’ ends before aligning to the *Sus scrofa* 10.2.69 reference genome. Additional trimming 6bp from the 5’ ends of cDNA reads was done to prevent misidentification of DNA RNA mismatches due to artifacts associated with the use of random hexamers during cDNA library preparation [17]. When conducting a search for RNA editing candidates with RNA-seq, strand-specific RNA-seq libraries can be utilized to account for the strandedness of each transcript, thereby enabling A-to-G DNA-to-RNA mismatches to be distinguished from T-to-C DNA-to-RNA mismatches. In order to utilize our strand-specific cDNA alignments for variant calling while preserving the strandedness of each alignment to distinguish A-to-G from T-to-C mismatches, plus-strand alignments were separated from minus-strand alignments for each cDNA sample. From all genomic DNA and cDNA alignments, we extracted those reads that had only 1 recorded alignment in order to optimize our chances that genomic DNA and cDNA reads arising from the same locus map to the same location. Joint variant calling using SAMTools [18] was performed, combining genomic DNA alignments with cDNA plus-strand alignments from each tissue. This was repeated for all cDNA minus-strand alignments. Both resulting VCF files were analyzed using editTools, an in-house R package made to efficiently scan VCF files for DNA RNA mismatches using C++ source code. editTools was developed to implement RNA editing detection within the R framework and to provide visualization tools; editTools was used to generate all figures in this manuscript pertaining to sequencing data. Default editTools parameters were used, in which a mismatch was considered a candidate RNA editing site if at a particular locus 1) the genotype is homozygous according to 95% of the DNA reads, 2) at least 10 reads were used to determine the genotype, 3) neither genomic DNA nor cDNA samples are indels, 4) at least 5 cDNA reads from the same tissue differ from the genotype call, and 5) these cDNA reads must have a Phred-scaled strand-bias P-value of 20 or less. Specific thresholds for DNA and cDNA sequencing depths were determined according to a previous study that profiled the rhesus macaque editome from a single animal [8]. Using this approach, we identified a total of 6410 A-to-G mismatch events representing 75% of all mismatches found (8550 total mismatches; Fig. 1). When we restrict our search to known swine repetitive sequences, 5993 out of 6410 A-to-G mismatches are retained, representing 93.8% of all mismatches in repetitive regions. Of the remaining mismatches in repetitive regions, 4.1% are T-to-C. It is not surprising that T-to-C mismatches are the second most common since T-to-C artifacts could arise if at a true A-to-G editing site, plus-strand alignments were incorrectly identified as minus-strand alignments or vice versa.

**Fig. 1.**
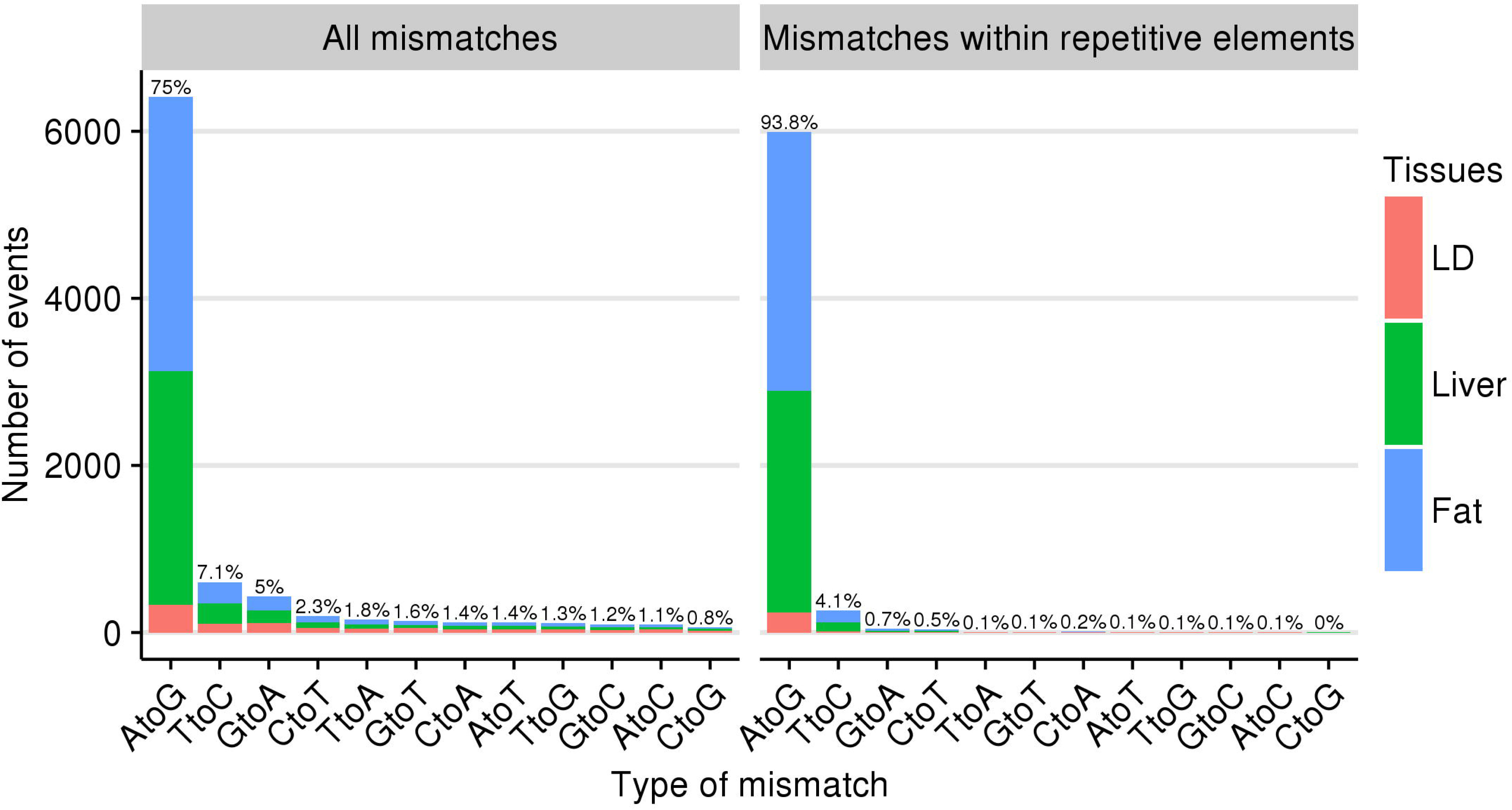
DNA to RNA mismatch counts. Comparing all mismatches found transcriptome wide (Left) to those within the body of a repetitive element (Right). Percentages shown are out of all mismatches found in each category.

### Tissue differences

To understand differences in candidate RNA editing sites between tissues, canonical A-to-G mismatches were aligned across tissues if they were detected at the same physical position and on the same strand. The number of candidate RNA editing events was fewer in LD compared to liver or fat (Fig. 1), consistent with lower RNA editing activity in muscle compared to other tissues for rhesus macaque [8]. Despite candidate RNA editing sites showing strong tissue specificity, a total of 144 A-to-G mismatches were found to be common among all three tissues, whereas 748 were found to be common between liver and fat (Fig. 2).

**Fig. 2.**
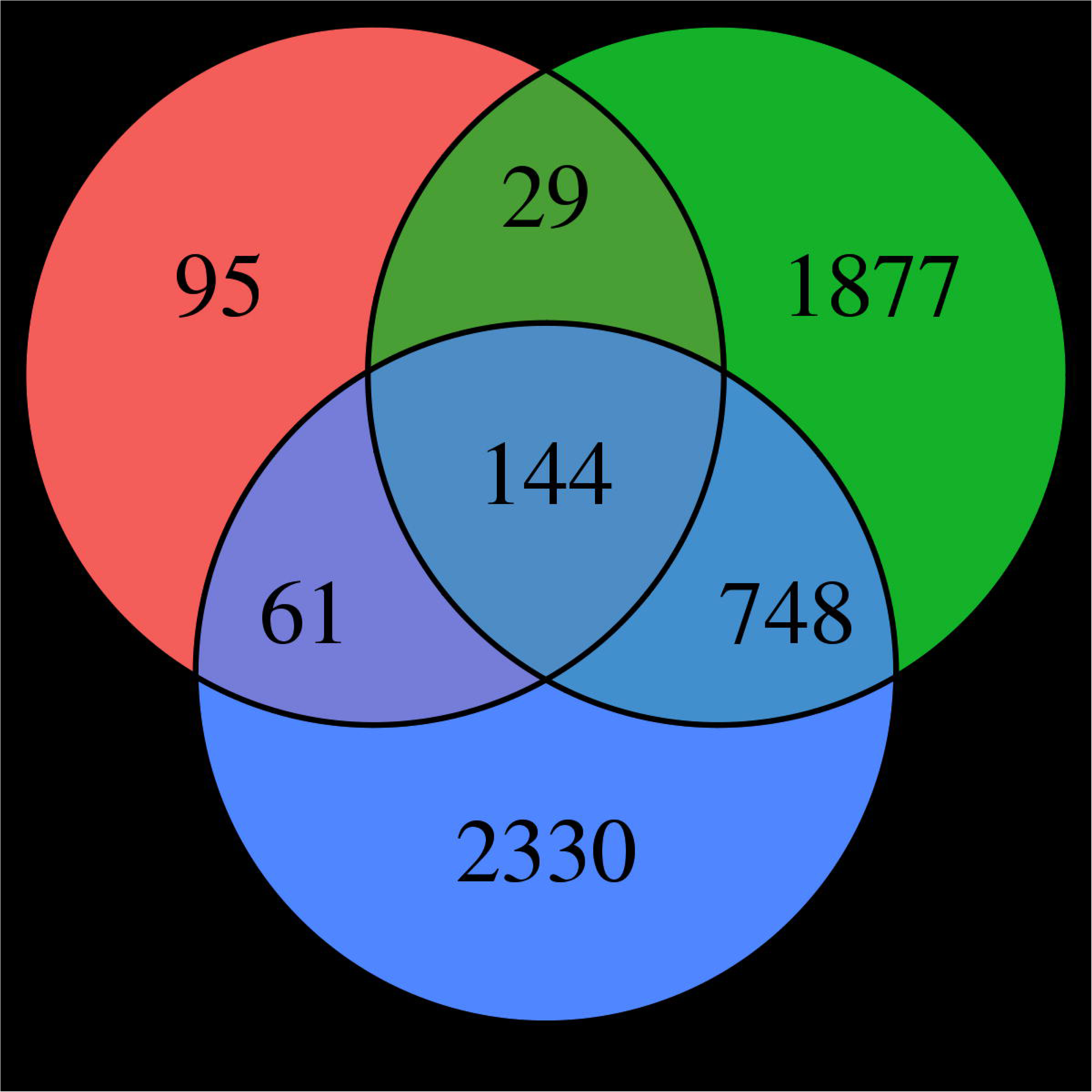
Shared A-to-G mismatches between tissues. A mismatch between two or more tissues was considered shared if it occurred at the same physical position and on the same strand.

One factor that may contribute to tissue specificity of RNA editing is differential expression of ADAR [19]. Using RNA samples from 33 additional pigs, a quantitative real-time PCR assay was used to infer ADAR transcript abundance differences between liver, subcutaneous fat, and LD muscle (Fig. 3). Average ADAR expression was determined to be significantly lower in LD muscle tissue than in either fat (p < 0.0003) or liver (p < 0.00001) tissues, suggesting that differential ADAR expression may contribute to differences in candidate RNA editing sites between tissues.

**Fig. 3.**
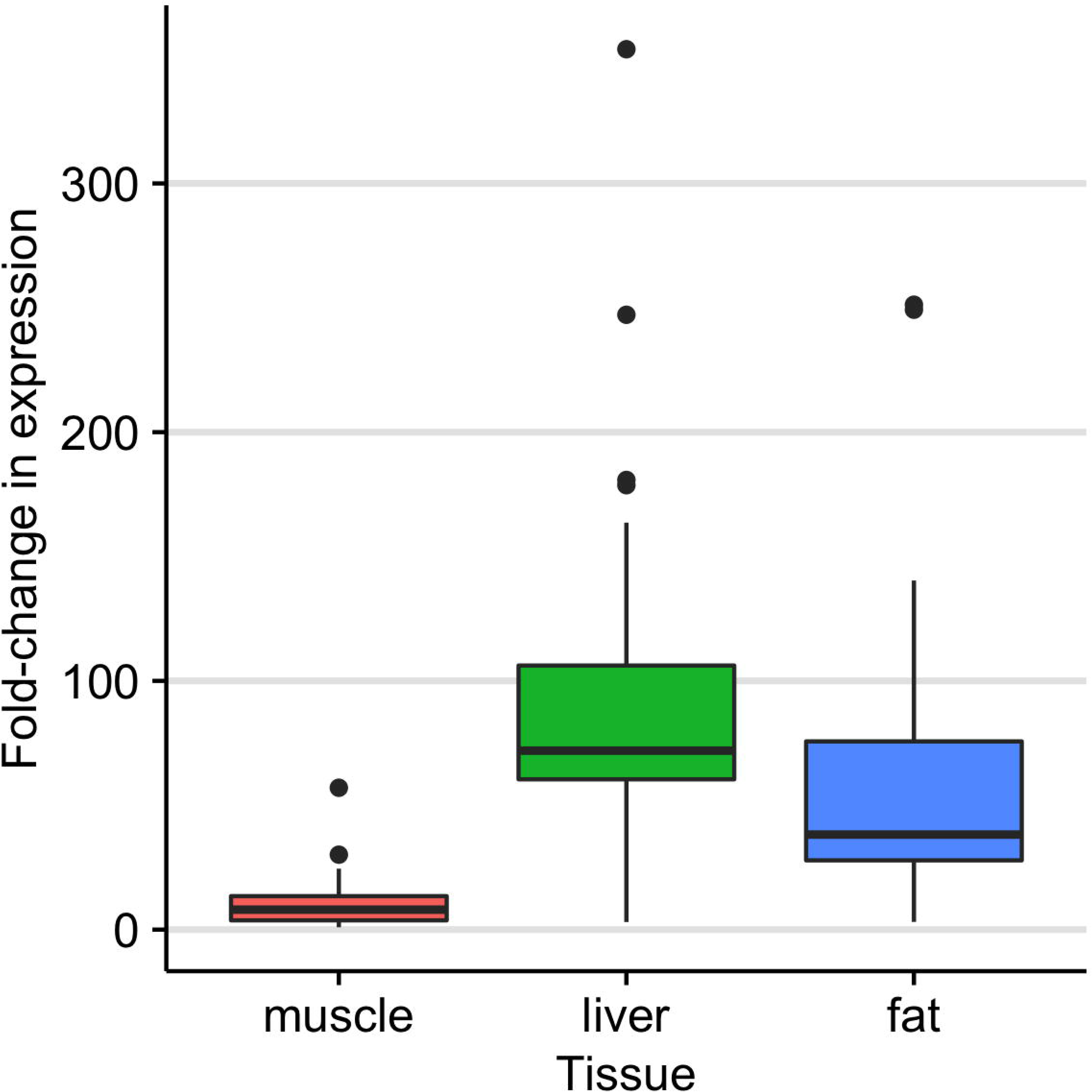
Relative ADAR transcript abundance between tissues. Expression was measured relative to the LD muscle sample used for sequencing. Using a one-way ANOVA, a significant effect of tissue on ADAR expression was detected (p < 0.0001). Pairwise comparisons of tissue means using Tukey HSD shows significant differences in ADAR expression between LD and liver (p < 0.00001) and between LD and fat (p < 0.003), but no significant difference between fat and liver (p = 0.0505563).

### Controlling for errors due to mapping quality

After imposing such strict restrictions as excluding genomic DNA and cDNA reads that had more than one recorded alignment and trimming the ends of reads pre-alignment, we wanted to assess how well such measures protect against mapping errors, which are among the leading causes of RNA editing misidentification when using short reads [17, 20]. Mapping quality is a measurement that provides a probability that a read is misaligned, given its number of possible alignments and sum of base qualities for each alignment [21]. Knowing this, and under the assumption of no RNA editing, for each mismatch locus *i* we computed the probability of observing at least 5 “edited” reads given the cDNA sequencing depth *N_i_* and average sample mapping quality *MQ_i_*. Among all 8550 repetitive and non-repetitive mismatch positions, the maximal probability of observing at least 5 “edited” reads was ~ 6.772e-15 for a site with *N* = 13 and average *MQ* = 29. If Bonferroni correction is used then 0.05 / 189,638 = 6.23e-07 can be used as a threshold for transcriptome-wide significance, where 189,638 was the total number of queried cDNA positions with a sequencing depth of at least 5 cDNA reads that were at the location of homozygous loci in the genomic sequence. From this evidence we conclude that our pipeline sufficiently minimizes artifacts associated with mapping quality when using the *Sus scrofa* 10.2.69 assembly.

### Pig editome functional implications

Little is known about the average effect of RNA editing transcriptome wide. For humans, one prevailing hypothesis is that the exonization of Alu SINE elements is controlled in part by A-to-G editing. An instance of this mechanism has been demonstrated, where intronic A-to-G editing events contribute to alternative splicing of *nuclear prelamin A* so that an Alu element is included in an exon [12]. To explore the possibility that RNA editing in pigs targets introns to affect splicing, editTools was used to synthesize mismatch data with Variant Effect Predictor data to find the relative locations of each mismatch relative to annotated transcripts. Consistent with what has been found in humans [2], nearly half of all detected A-to-G mismatches are located in retained introns (Fig. 4). The remaining sites are concentrated in other non-coding regions including 3’ UTRs, intergenic, and gene proximal regions. While the majority of non-coding editing events in humans are attributed to the position and orientation of SINE elements within transcripts [10], coding RNA editing occurs rarely, usually outside repetitive elements but nevertheless site-specifically. It has been suggested that site-specificity of coding RNA editing events is facilitated by nearby SINE elements, which through their induction of long dsRNA regions, recruit ADAR in sufficient density to affect coding regions in close proximity [16]. From our data, only 49 pig A-to-G mismatches were found within coding regions and of those, 34 would result in a missense variant (Table 1). It can be noted that a number of amino acid changes resulting from verified macaque DNA RNA mismatches [8] can be found among our pig dataset – mismatches that control I/V in *COPA*, Y/C in *BLCAP*, I/V in *COG3*, K/R in *NEIL1*, and Q/R in *GRIA2*. Interestingly, Y/C recoding of *BLCAP* via RNA editing has been associated with hepatocellular carcinoma (HCC) in humans as HCC samples were shown to express edited *BLCAP* in significantly higher amounts than non-HCC samples [22]. Additionally, exon 6 K/R recoding of *NEIL1* by RNA editing was previously thought to be primate specific and attributed to the K/R site’s proximity to Alu dense regions [23], however we witness evidence of the same K/R recoding of exon 6 via an A-to-G editing event in pigs. If in fact SINE elements recruit ADAR to affect nearby coding regions, then our data suggest the remarkable conservation of *NEIL1* K/R recoding across genomes with entirely different SINE elements.

**Fig. 4.**
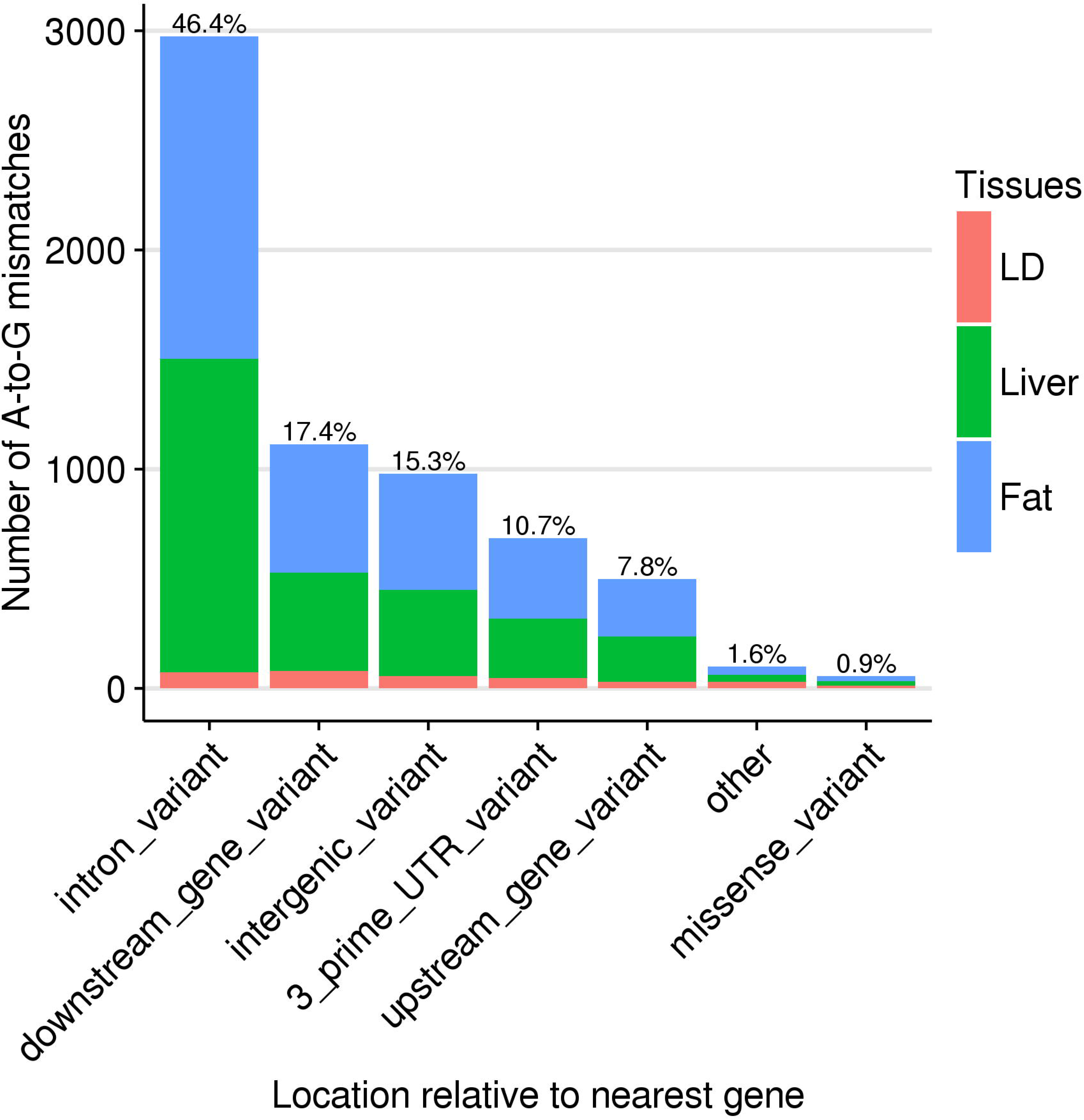
A-to-G mismatch locations relative to the nearest annotated gene. Percentages shown are out of all A-to-G mismatches.

**Table 1.**
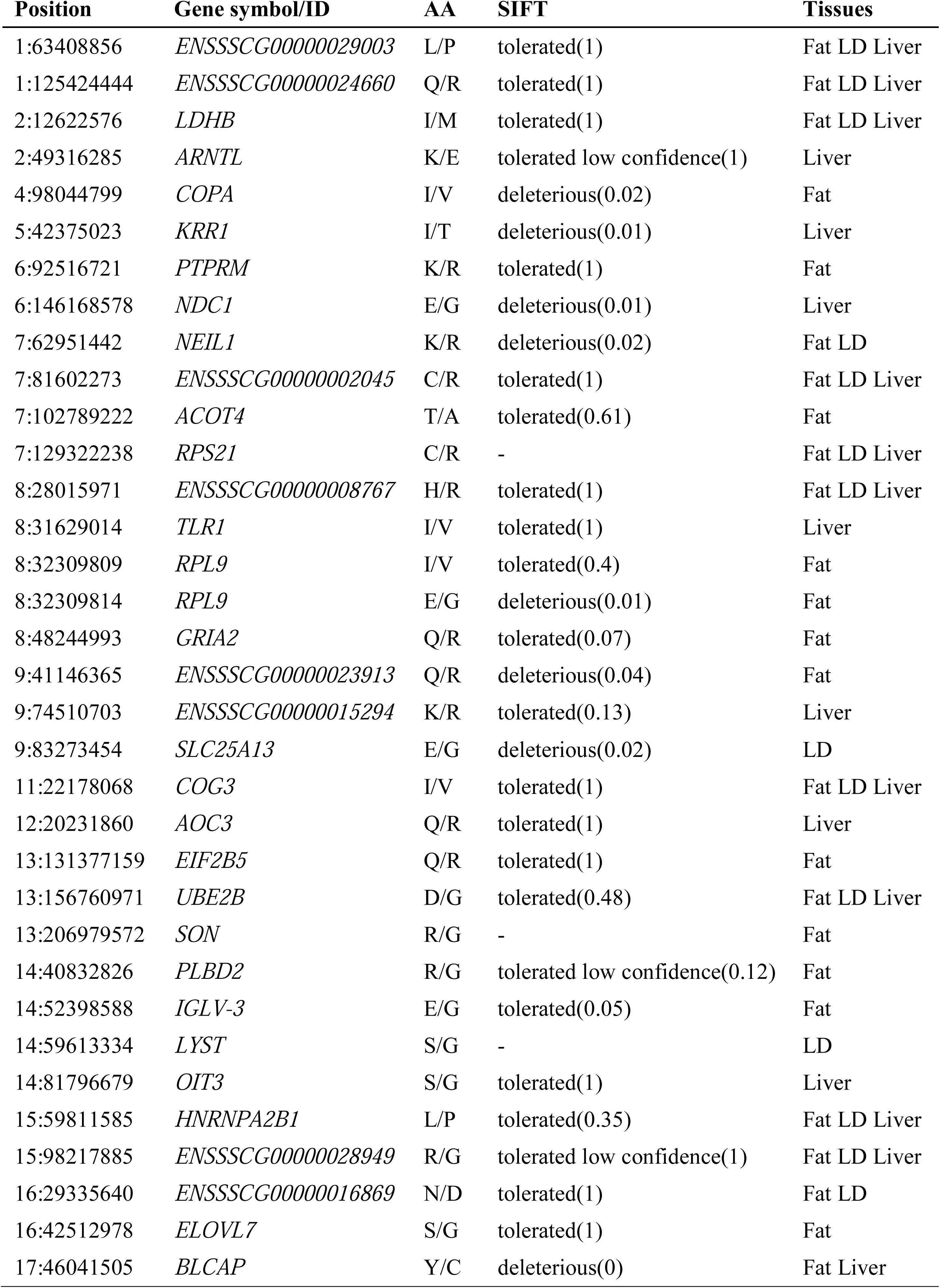
A-to-G mismatches resulting in amino acid changes.

### Pig editome association with pig-specific SINE elements

Since properties of the primate Alu element are suggested to influence RNA editing in both coding and non-coding regions, one of our primary interests was to determine which SINE elements in pigs are capable of attracting the majority of ADAR activity. Again using the functionality of editTools, we merged our mismatch data with data from RepeatMasker to determine which repetitive regions contain putative RNA editing sites. As mentioned previously, 5993 out of 6410 A-to-G mismatches are located within the body of a repetitive element. Upon closer inspection, 5715 of the 5993 are within pig SINE elements as opposed to LINE elements and others (Fig. 5A), although SINEs occupy just 11.4% of the swine genome, while LINEs occupy 17.5% [24]. Of the 5993 repetitive A-to-G mismatches, 58.8% are found within the Pre0_SS element, a SINE element of the PRE1 family (Fig. 5B). Little is known about Pre0_SS, but among all elements of the PRE1 family, Pre0_SS is most identical to the consensus PRE1 sequence. In many instances, Pre0_SS elements are > 99% identical to one another, indicating that it is currently actively transposing in pigs [25]. Additional members of the PRE1 family contain A-to-G mismatches, although at a much lower frequency than Pre0_SS.

**Fig. 5.**
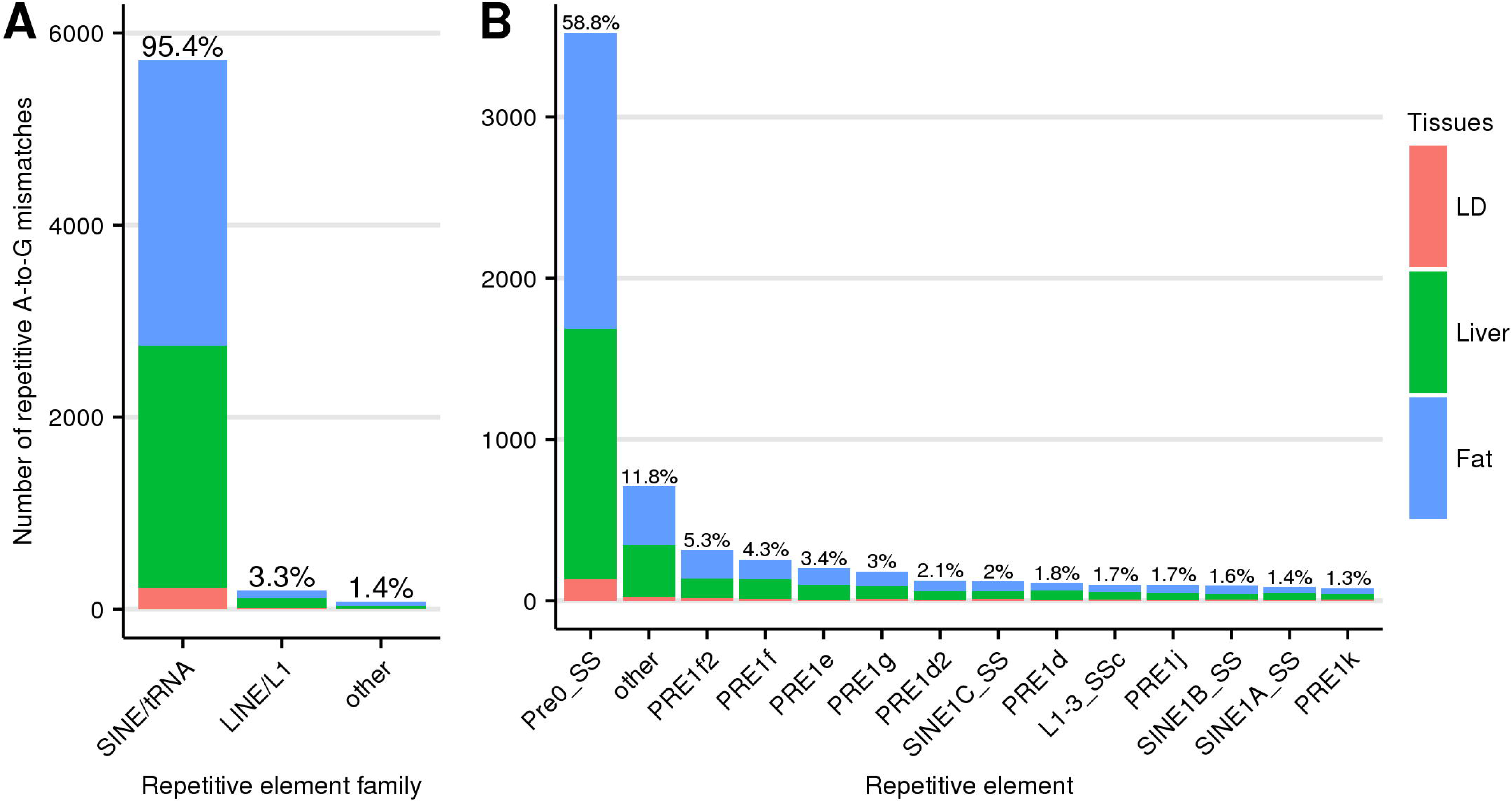
Distribution of repetitive A-to-G mismatches. The distribution is shown across major repetitive element families (A) and further broken down into specific repetitive element types (B). Percentages shown are out of all repetitive A-to-G mismatches.

## Conclusions

While Alu elements enable substantial RNA editing among primate genomes, we show that non-Alu bearing genomes can also utilize RNA editing as a means to achieve a similar result. Our high-throughput scan suggests that pig transcriptomes are highly editable among PRE-1 SINE retrotransposons. PRE-1, an element derived from an ancestral tRNA, has similar features to the primate Alu, derived from an ancestral 7SL RNA; a copy number of 1x10^6, consensus length of 246bp, and very little diversity among such members as Pre0_SS. These features influence the secondary structure of the transcriptome, which in turn affect ADAR editable targets. Surprisingly, conservation of specific editing sites such as those in *NEIL1* and *BLCAP* appears evident between human and pigs. Therefore, we hypothesize that transcriptome secondary structure may be conserved among mammals enough to preserve particular RNA editing sites, and that SINE elements, regardless of origin, may conform to certain positions and orientations in order to allow conservation to occur.

By demonstrating that pig transcriptomes have potential to be highly edited, we propose that pigs may be a valuable model to understand the patterns of ADAR controlled RNA editing. Additionally, by shedding light on the pig editome, we can begin to understand the extent to which this phenomenon enhances pig genetic variation. Such sources of variation may one day provide valuable explanatory power for a variety of traits of interest to both biomedical and agricultural communities.

## Methods

### Sequence data

From Michigan State University’s pig resource population (MSUPRP), an F_2_ population resulting from crosses between 4 F_0_ Duroc sires and 15 F_0_ Pietrain dams [26], a single female animal was chosen for whole genome and transcriptome sequencing. Total RNA was extracted from subcutaneous fat, liver, and LD skeletal muscle using TRIzol, and a RIN greater than 7 was determined with the Agilent 2100 Bioanalyzer. cDNA libraries were made using the Illumina TruSeq Stranded mRNA Library Preparation Kit. Sequencing was performed using the Illumina HiSeq 2500 in Rapid Run mode with 150x2 paired-end reads. Base calling was done by Illumina’s Real Time Analysis v1.18.61 and the output was converted to FastQ format with Illumina’s Bcl2fastq v1.8.4. Genomic DNA was purified from white blood cells using the Invitrogen Purelink Genomic DNA Mini Kit and libraries were made using the Illumina TruSeq Nano DNA Library Preparation Kit HT. Sequencing of genomic DNA was done using the Illumina HiSeq 2500 in Rapid Run mode with 150x2 paired-end reads. Real Time Analysis v.1.17.21.3 and Bcl2fastq v1.8.4 were used for base calling and FastQ conversion, respectively. Read quality of both whole genome and RNA data was assessed using the FastQC program [27].

### Sequence preparation and mapping

DNA reads from whole genome sequencing were trimmed for quality at the 3’ end using Condetri v2.2 [28] with parameters: -sc=33 -minlen=75 and b=fq. Resulting mate 1, mate 2 and unpaired reads were mapped to *Sus Scrofa* 10.2.69 using Bowtie v2.2.1 [29] with parameters: -p 7 -X 1000. In order to filter out DNA reads that had more than one recorded alignment, alignments containing the “XS:i:<N>” tag, where N indicates the number of alternative alignments for a read, were removed. Strand specific cDNA sequencing reads from each tissue sample were trimmed with Condetri with parameters: -sc=33 -minlen=75 -pb=fq -cutfirst=6 -pb=fq. Resulting paired and unpaired cDNA reads were then mapped to *Sus Scrofa* 10.2.69 using TopHat v2.0.12 [30] with parameters: -p 7 –mate-inner-dist 400 –mate-std-dev 100 –library-type "fr-firststrand”. Filtering out cDNA reads that had more than one recorded alignment was done by selecting alignments with the “NH:i:1” tag, while separating plus strand transcript alignments from minus strand alignments was done by selecting alignments possessing the “XS:A:+” or “XS:A:-” tags, respectively. The resulting DNA and cDNA alignments are the “filtered” data used in downstream variant calling and mismatch detection.

### Variant calling and mismatch detection

We utilized variant calling software Samtools v1.0 and Bcftools v1.2 to jointly call variants among DNA and cDNA reads from plus strand transcripts using: samtools mpileup –f <reference_genome.fa> –C50 –E –Q25 –ug –t DP,DV,SP <DNA.bam> <liver_plusstrand.bam> <fat_plusstrand.bam> <LD_plusstrand.bam>, where <DNA.bam> includes all filtered DNA alignments, and <liver_plusstrand.bam>, <fat_plusstrand.bam>, and <LD_plusstrand.bam> are filtered cDNA alignments from plus strand transcripts. Likewise, DNA and cDNA reads from minus strand transcripts were processed similarly with: samtools mpileup –f <reference_genome.fa>–C50 –E –Q25 –ug –t DP,DV,SP<DNA.bam> <liver_minusstrand.bam> <fat_minusstrand.bam> <LD_minusstrand.bam>. Note that the parameter “–t DP,DV,SP” is required for downstream mismatch detection with editTools. Samtools output from each command was piped into bcftools with additional parameters: –O v –m –v. These steps produce two VCF files that are simultaneously processed with find_edits(), a function within editTools available at https://github.com/funkhou9/editTools. By default, find_edits() scans each variant site to search for candidate RNA editing sites according to the five criteria required for sufficient evidence (see Results and Discussion). Most figures in this report were generated using editTools plotting methods, which utilized the ggplot2 R package [31].

### Quantitative real-time PCR

Total RNA was isolated from liver, LD skeletal muscle and subcutaneous fat tissues from 34 MSUPRP pigs, including the pig chosen for sequencing, using TRIzol reagent (Ambion) according to the manufacturer’s instructions. Concentrations were measured using a NanoDrop spectrophotometer (Thermo Scientific), and quality and integrity were determined using an Agilent 2100 Bioanalyzer (Agilent Technologies, Inc.). Total RNA was reverse transcribed using random primers with the High Capacity cDNA Reverse Transcription Kit with RNase Inhibiter (Applied Biosystems) according to the manufacturer’s instructions. A pig ADAR Custom TaqMan Gene Expression assay was designed using the online Custom TaqMan Assay Design Tool (ThermoFisher Scientific). The assay was designed to span exons 2-3 of the pig ADAR gene (Accession No. NC_010446.4). Assays were performed in triplicate using 50 ng cDNA and the TaqMan Gene Expression Master Mix (20 μl final volume per reaction) in a StepOnePlus Real-Time PCR System (Applied Biosystems). Cycling conditions were 52°C for 2 min and 95°C for 10 min, followed by 40 cycles of 95°C for 15 s and 60°C for 1 min.

Relative expression values were obtained using the 2^-ΔΔ^CT method, with the muscle sample used for sequencing as a calibrator and Ubiquitin C as a reference gene (Applied Biosystems Assay No. Ss03374343_g1). Inference of differential ADAR expression was calculated by one-way ANOVA (main effect of tissue on ADAR expression), and Tukey HSD (pairwise comparisons of tissue means).

### Calculating probability of mapping error

The average phred-scaled mapping quality *MQ* across all samples at mismatch site *i* is provided by SAMTools output. From *MQ* we can compute the probability of mapping error *p* according to:

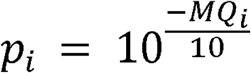

It follows that the probability of observing 5 “edited” reads at a homozygous site with a cDNA sequencing depth of *N* assuming no RNA editing can be modeled using the binomial distribution, where:

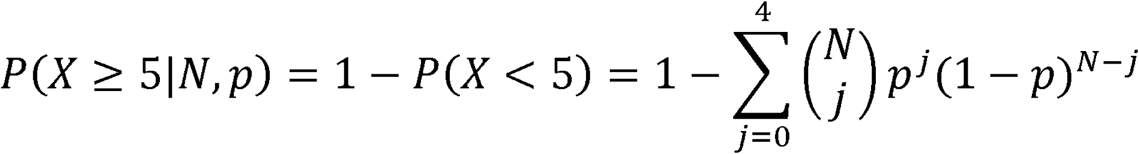

### Incorporating RepeatMasker and Variant Effect Predictor data using editTools

The editTools function add_repeatmask() was used to merge a mismatch data object (generated with find_edits()) with susScr3, a Repeatmasker dataset available for download at: http://www.repeatmasker.org/species/susScr.html. This function utilizes a binary search algorithm implemented in C++ to process large RepeatMasker files efficiently. The function write_vep() was used to generate Variant Effect Predictor input from a mismatch data object. The output of Variant Effect Predictor was merged with the mismatch data object using add_vep(). Additional documentation for find_edits(), write_vep(), add_vep(), add_repeatmask() is available within editTools v2.1.

## Abbreviations

ADAR: adenosine deaminase acting on RNA
LD: *longissimus dorsi*
LINE: long interspersed nuclear element
SINE: short interspersed nuclear element
UTR: untranslated region

## Declarations

### Ethics approval and consent to participate

Animal protocols were approved by the Michigan State University All University Committee on Animal Use and Care (AUF# 09/03-114-00).

### Consent for publication

Not applicable

### Availability of data and materials

Raw whole genome sequencing and RNA-seq data are accessible from the Sequence Read Archive, BioProject PRJNA354435.

### Authors’ contributions

Conceived and designed the study: CWE. Contributed samples from the MSU pig resource population: ROB, CWE, NER. Isolated RNA and DNA: NER. Developed software and analysis pipeline: SAF, JPS. Performed qPCR assays and analysis: DS, NER, SAF. Wrote the manuscript: SAF. All authors read and approved the final manuscript.

## Acknowledgements

Computing resources were provided by Michigan State University’s High Performance Computing Cluster. Sequencing was performed at the Michigan State University Research Technology Support Facility.

